# Structural Evolution of the Ancient Enzyme, Dissimilatory Sulfite Reductase

**DOI:** 10.1101/2021.12.28.474277

**Authors:** Daniel R. Colman, Gilles Labesse, G.V.T. Swapna, Johanna Stefanakis, Gaetano T. Montelione, Eric S. Boyd, Catherine A. Royer

## Abstract

Dissimilatory sulfite reductase is an ancient enzyme that has linked the global sulfur and carbon biogeochemical cycles since at least 3.47 Gya. While much has been learned about the phylogenetic distribution and diversity of DsrAB across environmental gradients, far less is known about the structural changes that occurred to maintain DsrAB function as the enzyme accompanied diversification of sulfate/sulfite reducing organisms (SRO) into new environments. Analyses of available crystal structures of DsrAB from *Archaeoglobus fulgidus* and *Desulfovibrio vulgaris*, representing early and late evolving lineages, respectively, show that certain features of DsrAB are structurally conserved, including active siro-heme binding motifs. Whether such structural features are conserved among DsrAB recovered from varied environments, including hot spring environments that host representatives of the earliest evolving SRO lineage (e.g., MV2-Eury), is not known. To begin to overcome these gaps in our understanding of the evolution of DsrAB, structural models from MV2.Eury were generated and evolutionary sequence co-variance analyses were conducted on a curated DsrAB database. Phylogenetically diverse DsrAB harbor many conserved functional residues including those that ligate active siro-heme(s). However, evolutionary co-variance analysis of monomeric DsrAB subunits revealed several False Positive Evolutionary Couplings (FPEC) that correspond to residues that have co-evolved despite being too spatially distant in the monomeric structure to allow for direct contact. One set of FPECs corresponds to residues that form a structural path between the two active siro-heme moieties across the interface between heterodimers, suggesting the potential for allostery or electron transfer within the enzyme complex. Other FPECs correspond to structural loops and gaps that may have been selected to stabilize enzyme function in different environments. These structural bioinformatics results suggest that DsrAB has maintained allosteric communication pathways between subunits as SRO diversified into new environments. The observations outlined here provide a framework for future biochemical and structural analyses of DsrAB to examine potential allosteric control of this enzyme.

## Introduction

Between 12-29% of the organic carbon that is delivered to the sea floor is mineralized by biological sulfate (SO_4_^2-^)/bisulfite (HSO_3_^-^) reduction (1). As such, SO_4_^2-^/HSO_3_^-^ reducing organisms (SRO) play substantial roles in the global sulfur and carbon cycles, both today (2, 3) and in the geologic past (4–6). Dissimilatory reduction of SO_3_^-^/HSO_3_^-^ to hydrogen sulfide (H2S) in most SRO is catalyzed by the enzyme dissimilatory sulfite reductase (Dsr) in a reaction that requires six electrons: SO_3_^2-^ + 6e^-^ + 8H^+^ ➔ H_2_S + 3 H_2_O (7). Based on fractionation of sulfur isotopes preserved in sulfide and sulfate minerals in rocks, dissimilatory SO_4_^2-^/HSO_3_^-^ reduction (presumably via a Dsr-like enzyme) is thought to have evolved as early as 3.47 billion years ago (6). The earliest evolving SRO may have reduced SO_3_^-^/HSO_3_^-^ (8–12), that would have been readily produced through solvation of sulfur dioxide (SO_2_) released into the biosphere via widespread volcanism on early Earth (13). Subsequently, the ability for SRO to use SO_4_^2-^ as an additional electron acceptor likely occurred in response to the gradual oxidation of Earth, culminating in the Great Oxidation Event (GOE) that occurred ~ 2.3 billion years ago (14). Preceding the GOE for several hundred million years, sustained production of O_2_ allowed for oxidative weathering of continental sulfides that led to the release of SO_4_^2-^ to oceans (15). At the same time, sustained production of O_2_ would have led to a decrease in the availability of HSO_3_^-^, since it is unstable in the presence of strong oxidants like O_2_ and Fe(III) (16), both of which became more abundant on an oxygenated Earth. The combination of decreased availability of HSO_3_^-^ and increased availability of SO_4_^2-^ may have represented the selective pressure to recruit ATP sulfurylase (Sat) that catalyzes the ATP-dependent activation of SO_4_^2-^ to adenosine 5’-phosphosulfate (APS), and APS reductase (AprAB) that reduces APS to HSO_3_^-^, thereby allowing for the use of SO_4_^2-^ as an oxidant. In potential support of this model, the reduction of SO_4_^2-^ to HSO_3_^-^ is an endergonic process (requires ATP), whereas HSO_3_^-^ reduction is exergonic and is the major energy conserving step during SO_4_^2-^ reduction (17). Further evidence in support of HSO_3_^-^ reduction preceding SO_4_^2-^ reduction comes from physiological studies that reveal higher growth yields in model SRO when grown with HSO_3_^-^ relative to those grown with SO_4_^2-^ (18, 19). As such, the use of SO_4_^2-^, which imparts an additional energetic burden on SRO, appears to be an adaptation to allow for respiration of an oxidant, SO_4_^2-^, that was much more widely available later in Earth history.

Extant Dsr enzymes are hetero-tetrameric and composed of two highly homologous A and B subunits thought to have evolved from gene duplication (20). Electrons for HSO_3_^-^ reduction derived from small organic molecules *(i.e.*, lactate) or H2 are thought to be transferred to DsrAB via a third labile subunit, DsrC, through two C-terminal cysteine residues (7). Thousands of DsrAB sequences have been generated from cultivars and from environmental amplicon-based or metagenomic surveys that have been used to characterize the ecology of SRO and/or to reconstruct their evolutionary histories (21–24). These studies have revealed that SRO inhabit a broad range of habitat types, including subsurface, hydrothermal, soil, and freshwater/marine sediment environments. Moreover, DsrAB are much more widespread throughout archaeal and bacterial lineages than suggested from cultivars only a decade ago (23, 24). In particular, metagenomic surveys have significantly expanded the known taxonomic and genomic backgrounds where DsrAB are found, although many of the taxa harboring DsrAB are uncultured and thus, the function of DsrAB in these organisms is inferred from closely related cultivars or based on phylogenetic clustering among defined DsrAB groups (23, 24).

The recovery of SRO and their corresponding Dsr sequences from environments that are subject to extremes of pressure, temperature, salt concentration, and pH indicate that the organisms harboring Dsr have diversified to function under diverse physiological conditions. For example, a novel DsrAB-encoding euryarchaeote (within the Diaforarchaea/Thermoplasmatota group) was recently discovered in moderately acidic (pH range of ~3.0 to 5.4), high temperature (~50°C to 75°C) springs in Yellowstone National Park (YNP) through metagenomic sequencing (11). Several nearly complete genomes were recovered across multiple springs and years that were representative of these organisms, and none encoded Sat or AprAB, consistent with the ability of these organisms to grow with HSO_3_^-^, but not SO_4_^2-^(11). Phylogenetic analyses suggest these euryarchaeote DsrABs to belong to the earliest evolving lineage that also includes sequences belonging to other thermophilic Archaea largely found in hydrothermal vents or hot springs, and that which are generally inferred to respire HSO_3_^-^ (but not necessarily SO_4_^2-^). These results add credence to the notion that Dsr evolved to allow for the reduction of HSO_3_^-^ and later diversified to allow for the reduction of SO_4_^2-^ in habitats characterized by more modest environmental conditions. However, while it is clear that DsrAB has substantively diverged at the primary sequence level, it is unclear if this divergence translates to structural variation and whether conserved structural features of DsrAB exist that are invariant to change, irrespective of environmental conditions.

To begin to assess whether structural changes have occurred throughout the evolutionary history of Dsr, we modeled the structural characteristics of early-branching archaeal Dsr (i.e., from the newly characterized euryarchaotes described above) and compared these to those from an early-diverging group of Dsr (i.e., from the Archaeoglobales) and from a later-evolving group of Dsr (i.e., from *Desulfovibrio* and other Deltaproteobacteria). Sequence co-evolution (co-variance) analysis can provide information about residue-residue contacts and on functional coupling between distant sites (25–28). Sequence conservation and co-variance analysis of DsrAB sequences in the context of representative three-dimensional structures and models across the family was used to identify functionally-critical direct interactions, as well as longer-range potential functional couplings consistent with an intersubunit allosteric network, reminiscent of the negative cooperativity in the Mo-Fe nitrogenase (29). Observations of structural evolution are discussed in the context of environmental- and taxonomic-level adaptations that are likely to have taken place during the evolution of Dsr and the organisms that encode these enzymes.

## Methods

### DsrAB Database Generation and Phylogenetic Analyses

To evaluate the structural properties that confer Dsr function over broad geochemical space, an existing DsrAB database was curated from cultivar and environmental genomes (23). The original database was constructed to include DsrAB from genomes of cultivars, targeted PCR-based surveys, and metagenomic data. Thus, many of the sequences were incomplete, potentially confounding structural modeling calculations. Therefore, sequence alignments and annotations were used to curate the database to comprise only full length DsrA and DsrB sequences. Specifically, sequences demarcated as “partial” and those that were likely obtained via PCR-based methods were removed without further consideration. Next, individual alignments of DsrA and DsrB were performed using Clustal Omega (30), guided with primary sequences of DsrAB from *D. vulgaris* and *A. fulgidus*. Sequences that were substantially truncated relative to model DsrAB, including those without start codons, were then removed, resulting in a total of 274 full-length DsrAB sequences. The database will be made available upon request from the authors. Phylogenetic analysis of DsrAB sequences was conducted, as previously described (11), and associated metadata was mapped to the DsrAB phylogeny using environment of sequence origin (from the original database publication), in addition to taxonomic information (either from the original database publication or via BLASTp searches of DsrA subunits against the NCBI nr database). DsrAB sequence homologs were subjected to structural alignment using the PROMALS3D multiple sequence and structure alignment server (31). Structures from *D. vulgaris* (2V4J; (32)) and *A. fulgidus* (3MMC; (20)) served as threads.

### MV2-Eury Structural Homology Modeling

Comparative modeling of MV2-Eury Dsr A, B, and C subunits was performed after careful refinement of a structural alignment using ViTO (33) prior to model building using MODELLER (34) as a heterohexamer. The model was based on two templates PDB 3ORI1 (35) and PDB 3MM5 (36). The structure of the MV2-Eury heterotetramer was also modeled with AlphaFold2 (AF2) Multimer (37) from Deepmind, Inc. that is enabled for modeling of complexes, installed on the Rensselaer Polytechnic Institute’s Center for Computer Innovation NPL cpu/gpu cluster (https://secure.cci.rpi.edu/wiki/clusters/NPL_Cluster/) providing sufficient random-access memory and accessibility to handle these relatively-large multi-chain protein complexes.

### Evolutionary Co-variance and Anisotropic Network Analysis

Evolutionary covariance (EC) analysis (25, 26, 38) was performed using the EV Couplings server (https://evcouplings.org/ (39)) to identify instances of False Positive Evolutionary Covariance (FPEC), defined as significant covariance between residues that are too distant (>20 Å) to be in direct contact within the monomeric subunits. Given the strong homology between DsrA and DsrB subunits that are suspected to have originated from gene duplication, it was not possible to use the EV Complex Couplings option to identify co-varying residues at subunit interfaces in the oligomers. Instead, EC analysis was carried out using the monomeric sequences of the DsrA and DsrB subunits of *A. fulgidus*, *D. vulgaris*, and MV2-Eury separately as input. The values of Neff (number of non-redundant sequences in the alignment normalized to the number of residues) was between 1.16 and 2.84, depending upon the query, with the number of non-redundant sequences was between 481 and 2786. Queries were made by separately submitting the A and B subunits from the *A. fulgidus* and MV2-Eury sequences. Recovered coupling probabilities were between 80 and 94%. To model the possible dynamic fluctuations that might be associated with the putative allosteric pathway that was identified through EC analysis, we carried out Anisotropic Network Modeling (ANM) (40) using the webserver from the Bahar group (http://anm.csb.pitt.edu/cgi-bin/anm2/anm2.cgi; (41)).

## Results

### Phylogenetic Reconstruction of DsrAB

The curated database of full length DsrAB sequences (n=274) was used to construct a concatenated DsrA and DsrB phylogeny, as described previously (11), that was congruent with previous phylogenetic reconstructions of DsrAB (11, 23, 24). As previously documented, recently discovered Euryarchaeote DsrAB formed a group with those from Crenarchaeota, that together formed a basal-branching group among all DsrAB (Figure 1). All organisms within this first group were recovered from hydrothermal environments (largely hot springs) and their DsrAB are inferred to be involved in HSO_3_^-^ or SO_4_^2-^ reduction (11). An additional basal-branching second group comprised DsrAB recovered from uncultured metagenome-assembled-genomes (MAGs) from various organisms and environments, along with duplicate DsrAB copies within *Moorella* sp. genomes. The remainder of DsrAB comprised a large group inclusive of both reductive- and oxidative-type DsrAB that primarily includes bacterial DsrAB. Within the “bacterial” group, homologs from Archaeoglobales (Archaea) form a relatively early-evolving group, although the presence of DsrAB in Archaeoglobales is thought to derive from a horizontal gene transfer (HGT) event from a bacterial donor (22, 23). Lastly, the *D. vulgaris* homologs were present within a large cluster comprising homologs from other putative and characterized SO_4_^2-^ reducing Deltaproteobacteria.

**Figure 1.**
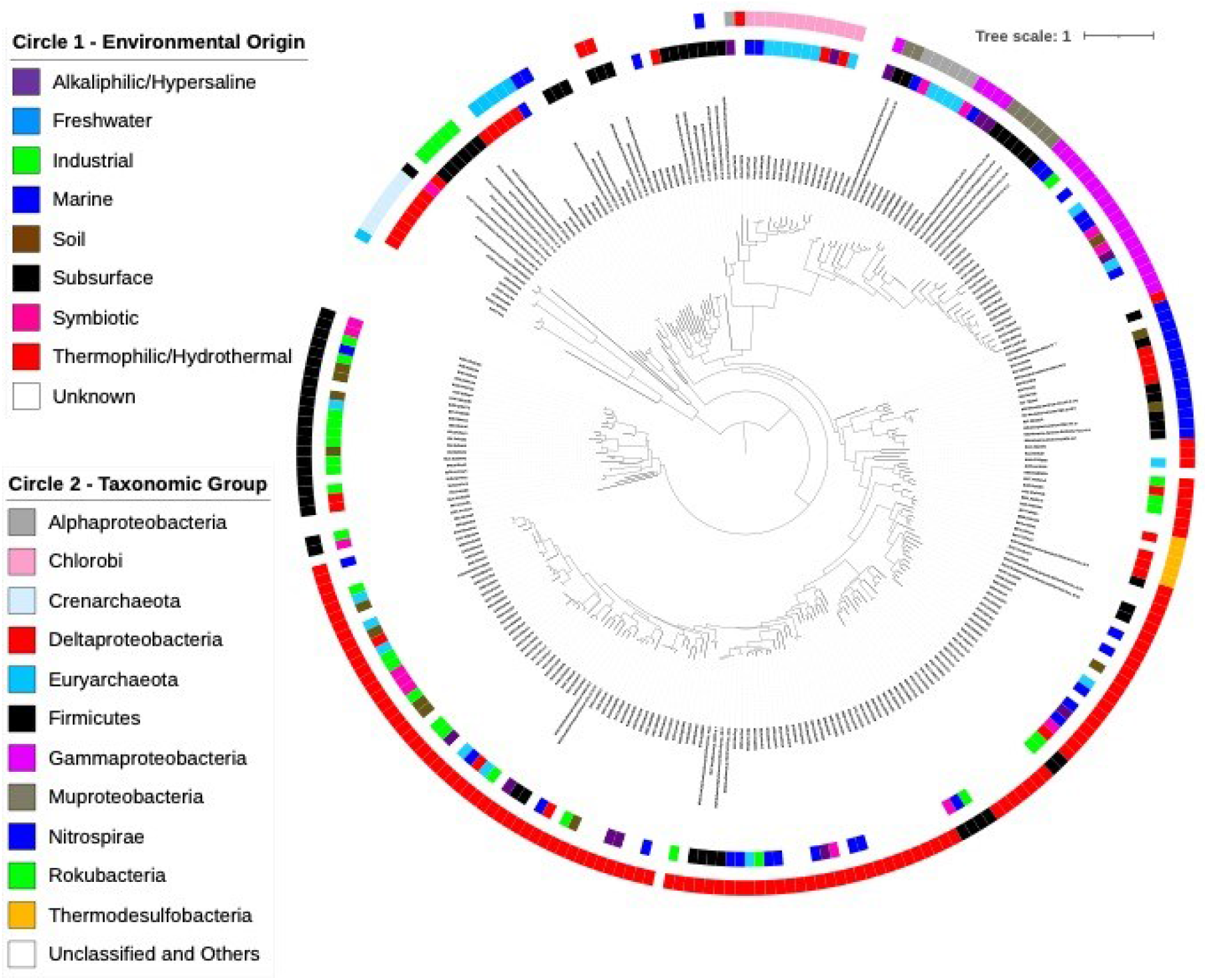
DsrAB phylogenetic reconstruction showing the taxonomic affiliation of Dsr-hosting organisms and their environmental origin. The Maximum Likelihood reconstruction was conducted on a concatenated alignment of DsrA and DsrB subunits from a curated database comprising the previously recognized primary homolog groups. The scale bar shows the expected substitutions per site. The environmental origin of Dsr homologs is shown based on available metadata associated with the previously published database (Muller et al. 2015) or from metadata associated with MAGs from metagenomes. Taxonomic classification is given based on information in the aforementioned database or by >80% amino acid homology to Dsr from cultivars genomes with taxonomic annotation.

Mapping of taxonomic information on to the DsrAB phylogenetic tree revealed general concordance of DsrAB clades with their respective taxonomic groups, consistent with previous analyses (23, 24). This indicates that DsrAB are generally vertically inherited, although several exceptions to this rule are evident including the example of Archaeoglobales above. The mapping also revealed the broad range of ecological contexts for SRO and their DsrAB. As documented previously (11), DsrAB from organisms with subsurface and hydrothermal environmental origins are particularly prominent near the root of the tree, suggesting that the earliest DsrAB may derive from oxidant limited and/or high temperature environments (42). However, general patterns of ecological distributions beyond these early groups were not readily apparent. This is likely attributable to the coarseness by which these original environmental designations were assigned (i.e., by site of isolation/sequence generation).

### Analysis of Available Dsr Crystal Structures

Three dimensional crystal structures have been solved for DsrAB enzymes from *Archaeoglobus fulgidus* (Figure 2A-B) (20, 36), *Desulfovibrio vulgaris* (32) (Figure 2C-D), *Desulfovibrio gigas* (35), and *Desulfomicrobium norvegicum* (43) at 2, 2.1, 1.76 and 2.5 Å resolution, respectively. The A and B subunits consist of three domains, two of which are structurally similar. The third is a ferredoxin-like domain, thought to have been inserted between two beta strands of domain two after the gene duplication event (20). Each A/B heterodimer in DsrAB harbors two sirohemes (or one siroheme and a sirohydrochlorin moiety, representing a siroheme without metal cofactors) and four [4Fe-4S] clusters that are presumably involved in electron transfer to HSO_3_^-^ (present in the *D. vulgaris* structure Figure 2B, D). A third, cysteine disulfide-containing labile subunit, DsrC, was purified and crystalized covalently bound to the heterotetramer in the *D. vulgaris* structure (32) (grey and purple in Figure 2B, D) via one of the reduced cysteine residues. In the most recently proposed model of the HSO_3_^-^ reduction reaction cycle, DsrC in reduced form binds to a S^II^ intermediate at the active site in DsrAB, forming a S^I^ containing hetero-disulfide. Hydrogen sulfide is then produced via an S^0^ -containing protein tri-sulfide intermediate, implicating both cysteine residues in the DsrC C-terminus (7). The final four electrons required for this latter reaction are proposed to emanate from the menaquinone pool, likely implicating the membrane DsrMK(JOP) complex (44), thus coupling HSO_3_^-^ reduction to proton translocation and energy conservation (7).

**Figure 2.**
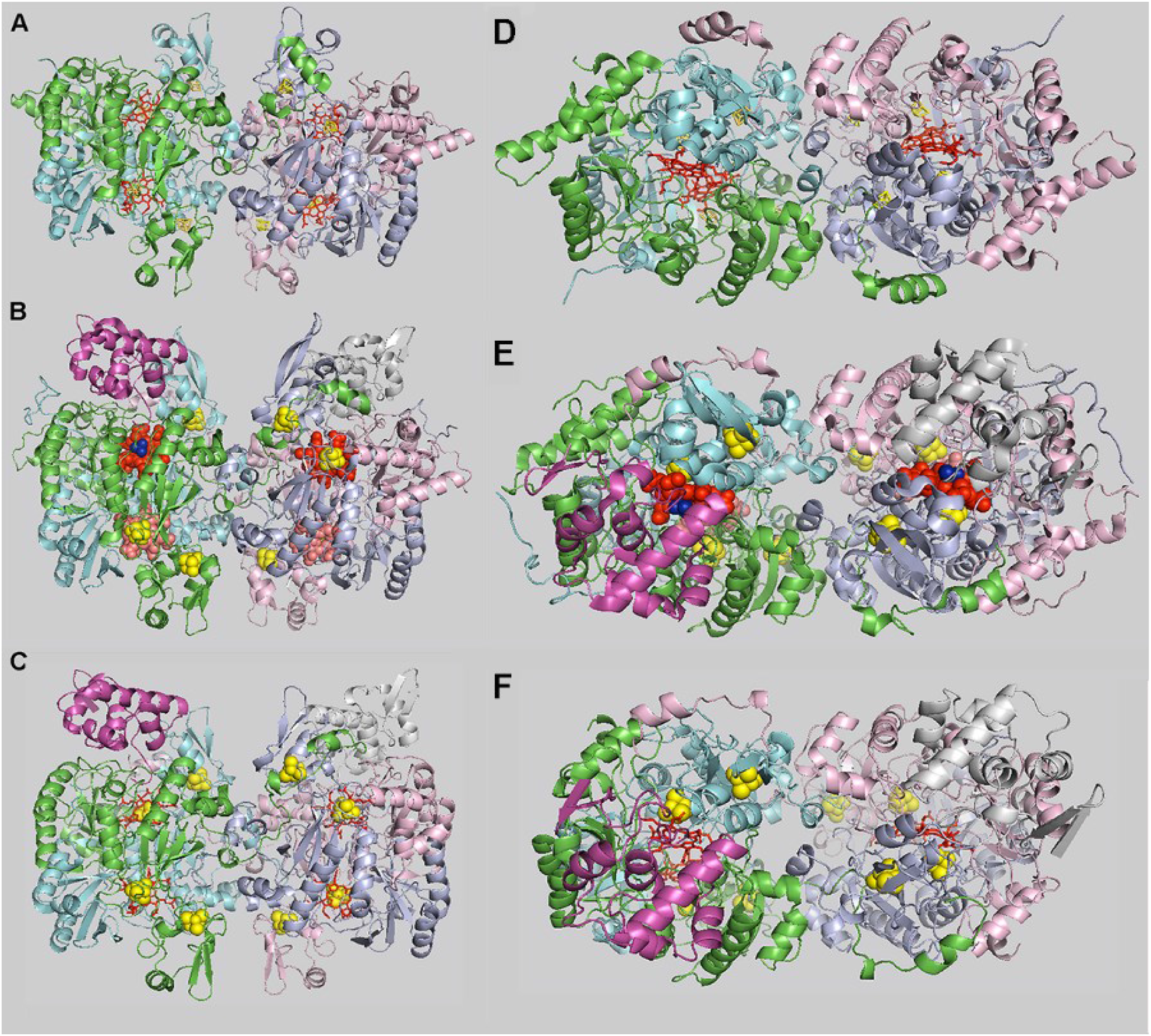
Available Crystal Structures of Dsr and a Homology Model of an Early Evolving Dsr From a Sulfite Reducing Euryarchaeote (MV2). A-C) side view; D-F) top view; A and D) *Archaeoglobus. fulgidus* heterotetramer - pdb id: 3mmc, B and E) *Desulfovibrio. vulgaris* hetero-hexamer – pdb id: 2v4j, C and F) homology model of MV2-Eury hetero-hexamer. Chain colors are: A1 (green), B1 (cyan), A2 (light pink), B2 (light blue), C1 and C2 where present (purple and white, respectively). Sirohemes (red), siro-hydrochlorine (salmon), and Fe-S centers (yellow) are sticks or spheres. The sulfite substrate is shown in the *D. vulgaris* structure (B) in dark blue spheres.

Within the DsrAB heterodimer, the A and B subunits are intricately intertwined in the manner of clasped hands. In addition, the C-terminus of the A subunit in one heterodimer crosses over the heterodimer interface to interact extensively with both the A and B subunits of the other (Figure 2A). While crystallographic domain swapping has been demonstrated to be artefactual in many cases, the fact that all four available DsrAB structures (20, 32, 35, 43) exhibit this feature supports the notion that it is a fundamental feature of the DsrAB structure. Besides this crossover interaction, the central interface between heterodimers is quite open, with only limited contacts between two helices of the B subunits from each heterodimer (Figure 2 D, E). Interestingly, at this central interface between heterodimers, the position of the two helices from the B1/B2 subunits responsible for the contacts has been swapped between the *A. fuglidus* and *D. vulgaris* structures. All three of the other available structures of DsrAB (32, 35, 43) show the central interfacial helix positioning of the *D. vulgaris* structure shown in Figure 2D. Native gel electrophoresis followed by mass spectrometry measurements made on preparations of DsrABC from *D. vulgaris* and *D. norvegicum* revealed that the major oligomeric species were A2B2C2 and A2B2C heterohexamer and heteropentamer, the C subunit being somewhat labile (43). These observations reinforce the crystallographic results indicating that DsrAB is a heterotetramer.

It is thought that only two of the siroheme moieties of Dsr (one per heterodimer) support HSO_3_^-^ reduction (20). While the *A. fulgidus* heterotetramer (purified and crystalized anaerobically) comports four siroheme moieties, the *D. vulgaris* structure (purified aerobically) has only two, the other two being sirohydrochlorin groups lacking the iron. In this latter structure, the HSO_3_^-^ ion is found in interaction only with the sirohemes (Figure 2B, dark blue). In the *A. fulgidus* structure, access to the “bottom” siroheme is blocked by tryptophan B119 (Figure 3A), whereas in the *D. vulgaris* structure, no blocking residue is apparent near the sirohydrochlorin group (Figure 3B), the corresponding side chain of threonine B135 facing away from the heme.

**Figure 3.**
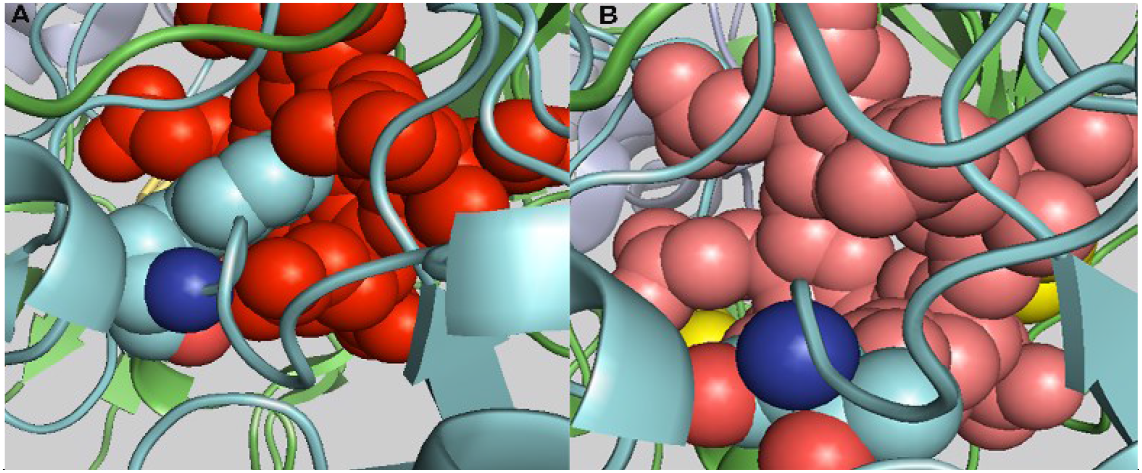
Close-up view of the “lower” prosthetic group in the structure of Dsr. A) *A. fulgidus* and B) *D. vulgaris* Chain colors are: A1 (green), B1 (cyan), Sirohemes (red), siro-hydrochlorine (salmon) and Fe-S centers (yellow) are sticks or spheres. Trp 119 and Thr 135 are shown in spheres and colored by element.

### Structural Homology Modeling of MV2-Eury DsrAB

The structural homology model of the MV2-Eury DsrABC sequences (Figure 1E, F), was based on the hetero-hexameric structure from *D. vulgaris* (32) due the poor quality of the electron density for the DsrAB structure from *A. fulgidus*. An obvious difference between the MV2-Eury model and the template is that the β-hairpin of the DsrB subunit of MV2-Eury that contacts DsrC (Figure 1E) is much shorter than in *D. vulgaris*, whereas the pseudo-symmetric β-hairpin structure in the MV2-Eury DsrA subunit near the inactive siroheme (Figure 1E, bottom, green & pink) is much longer. Examination of the sequences and differences between the MV2-Eury Dsr model and the *A. fulgidus* heterotetramer or the *D. vulgaris* hetero-hexamer structures suggested that much of the sequence variation among these three homolog types was found at subunit interfaces (Figure S1).

To complement the homology model in Figure 1, the MV2-Eury DsrAB heterotetramer was modeled also using AF2 Multimer (37). In this case the Fe-S centers and siroheme moieties are not present in the modeled structure. Three models were obtained, with the highest confidence (highest pLDDT scores) found for model 3 (Figure 4, Figure S2). As can be seen in the model colored according to the pLDDT score (Figure 4, Figure S2), the only regions of the complex that were poorly predicted in the models were in subunit A, residues 50-62 and 397-300. The first region interacts with the DsrC subunit (Figure 1B), while the second corresponds to four residues at the end of the first β-strand in a β-sheet adjacent to the A-subunit ferredoxin domain. Alignment of the MV2-Eury AF2 model with the homology model shows reasonable agreement (Figure 4), although the heterodimers in the AF2 model are rotated with respect to each other around the central axis of the heterodimer interface. Other notable differences are the orientations of the side chains at central interface between heterodimers (Figure S3) and a different orientation of the extended C-terminal A subunit helices that contact the opposing subunits. Note that the orientation of this helix is different in the *A. fulgidus* and *D. vulgaris* structures as well. The region with the low pLDDT scores (residues 50-62 in the A subunit) in the AF2 model exhibits a similar configuration to that in the homology model (Figure S3) but is rotated away from the center of the protein.

**Figure 4.**
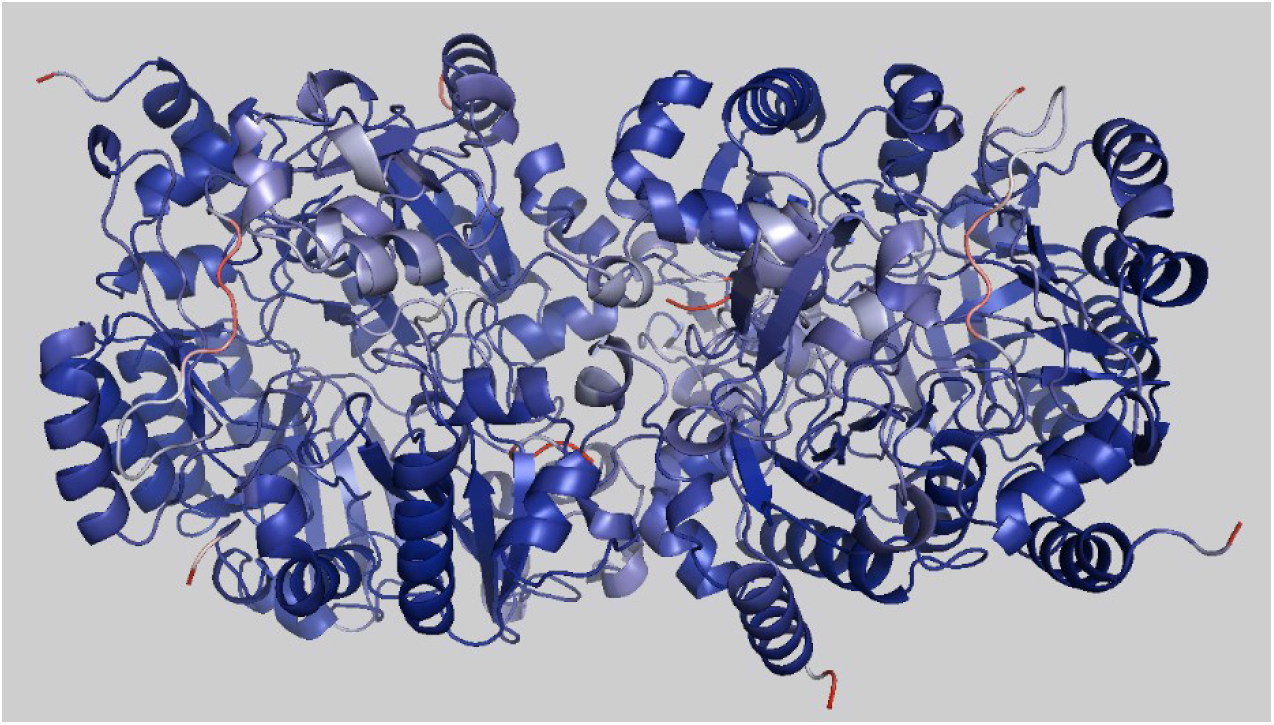
AlphaFold2 model of MV2.Eury DsrAB. The sequence is colored for pLDDT score (descried in the text), with dark blue corresponding to high confidence prediction (96%) and red to low confidence prediction (43%).The view is a top view of the structure as in Figure 1F. The light blue to red region on the center left and right correspond to residues 50-62 if the A subunit which interact with the DsrC electron donating subunit (Figure 1E, F). The upper left, bottom right and center red regions are the chain termini.

### Evolutionary Co-variance Analysis of DsrAB

Because many of the non-conserved residues in DsrAB are found at subunit interfaces, Evolutionary co-variance (EC) analysis on the EVCouplings server (as described in Materials and Methods) (39) was used to reveal how subunit contacts may have evolved with environment types or taxonomic groups that are distributed across the DsrAB phylogeny. We were particularly interested in False Positive Evolutionary Couplings (FPEC), as these often correspond to inter-subunit contacts within oligomers. They also can arise from long range allosteric pathways or dynamic structural heterogeneity (26), although these types of FPECs represent a small fraction of the total co-varying residues (45). Numerous FPECs were identified as evolutionarily co-varying residues that did not correspond to a contact in the 2D residue contact map of the monomer and were separated in space (within the monomers) by greater than 20 Å. Because the DsrA and DsrB subunits are highly homologous, both in sequence and in structure (Figure S4), both subunits were present in the multiple sequence alignments (MSAs) used in the EVCouplings analysis (38, 39). Moreover, because DsrAB complexes are hetero-tetramers, in principle, putative co-varying pairs of residues could correspond to eight different possible residue pairs (Res1A1-Res2A1, Res1B1-Res2B1, Res1A1-Res2B1, Res2A1-Res1B1, Res1A1-Res2A2, Res1A1-Res2B2, Res2A1-Res1B2, and Res1B1-Res2B2).

### Identification of a Potential Allosteric Pathway in DsrAB

Two sets of high co-variance probability (>80%) FPEC pairs (Table 1, bold) were observed using the *A. fulgidus* DsrA and DsrB subunits as queries in the EVCoupling calculations. Table 1 lists the homologous residues in the DsrB (or DsrA) subunits for these FPEC pairs for the three DsrAB structures in Figure 2, along with the FPEC distance and the coupling probability. Given the large distances between the putative co-varying residues in the monomers, and hence the false positive nature of the pair, the Res1A1-Res2A1 and Res1B1-Res2B1 co-variances are eliminated, leaving six remaining possible pairs of co-varying residues. Highlighting the four residues from the two A and two B subunits (eight residues in all) on the DsrAB structure (Figure 5A-C), it is particularly striking that three out of four of the possible implicated residues for each of the two sets of monomer FPEC pairs describe a path from the functional heme in one heterodimer to that in the other heterodimer. We refer to this pathway as the “heme road”. The fourth possible co-varying residue (Table 1) from each heterodimer, N222A1/2, is in a homologous structural position as N180B in the structural alignment of the A and B subunits (Figure S4), but in the heterotetramer N222A1/2 interacts with the “lower” inactive, structural siroheme at a distance of 20.5 Å from the nearest of the three other possible co-varying residues, whereas the homologous N180 of the DsrB subunit is proximal to the “upper” active siroheme and one of the nearby Fe-S centers, and constitutes the first residue in the heme road.

| 1. FPEC pairs for the heme road |  |  |  |  | Distance<br>in monomer<br>(Å) | Coupling<br>probability<br>(%) |
| --- | --- | --- | --- | --- | --- | --- |
| Organisms | Res1<br>DsrA | Res 2<br>DsrA | Res1<br>DsrB | Res2<br>DsrB |  |  |
| *A. fulgidus* | **<u>N393</u>** | **N222** | **<u>T351</u>** | *N180* | 33.51 | 90 |
| *D. vulgaris* | N410 | N223 | P368 | N191 |  |  |
| MV2-Eury | **N369** | **L204** | R337 | S167 | 33.84 | 80 |
| *A. fulgidus* | N392 | K240 | **I350** | **R197** | 23.62 | 94 |
| *D. vulgaris* | T409 | K241 | D367 | R208 |  |  |
| MV2-Eury | T368 | E222 | L336 | T184 |  |  |
| FPEC pairs resulting from EVCouplings analysis are shown in bold. All corresponding homologous residues in the *A. fulgidus*, *D. vulgaris* and MV2-Eury A and B chain sequences are shown. Distances are Cα-Cα distances. Bolded sequences correspond to FPEC residues from the queried subunits, italicized and underlined sequences to probable co-varying pairs (see text). | | | | | | |

**Figure 5.**
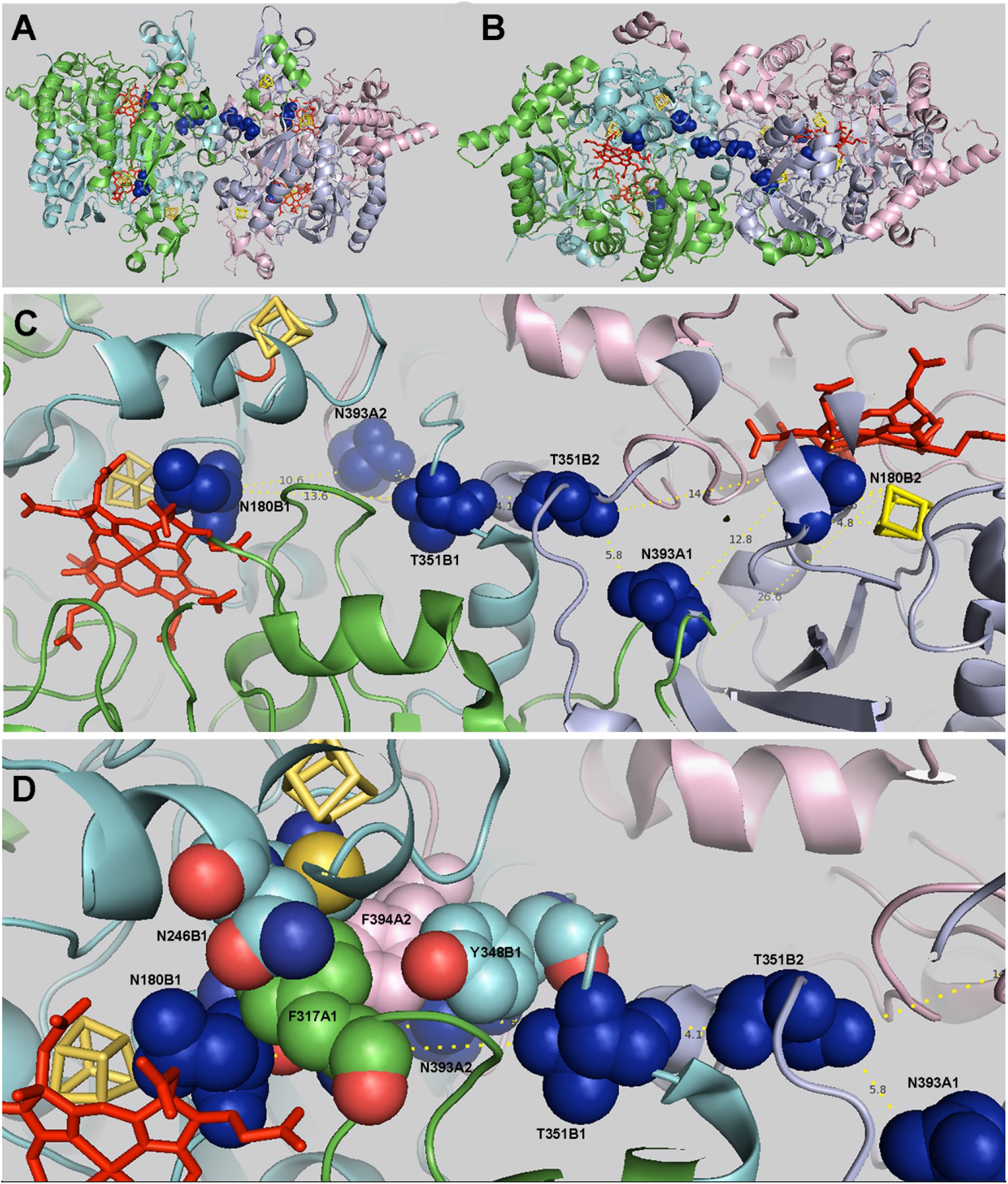
The heme road FPEC residues in the structure of DsrAB from *A. fulgidus*. A) Side view B) top view, C) slab of a zoom from the top view Distances between FPEC residues are shown in yellow. Chain colors are: A1 (green), B1 (cyan), A2 (light pink), B2 (light blue). Sirohemes (red sticks) and Fe-S centers (yellow) are also shown. Heme road FPEC residues (blue spheres) tracing a connection between functional hemes are from left to right N180B1, N393A2, T351B1, T351B2, N393A1 and N180B2. N222A1/2 (also blue spheres) interacts with the propionate group of the structural (non-functional) heme visible in (A) in a lower plane. D) Network of aromatic residues connecting the residues of the heme road (dark blue) for the left half of the heme road. Three aromatic residues, F317A1, F394A2 and Y348B1, directly connect the heme road residues. These interactions are stabilized from above by N246B1. N246B1 also contacts C244B1 that makes contact with the second Fe-S center distal to the siroheme. Two heme road residues T351B2 and N393A1 of the second heterodimer also appear on the right of the figure.

Residue N393 in the DsrA subunit is aligned to T351 in the DsrB subunit, but these occupy very different spatial positions (Figure S4). The N393 from the DsrA subunit is part of the C-terminal arm that extends to the other heterodimer (Figures 1, 5), such that N393A2 inserts itself into the DsrB1 subunit and vice-versa. This is the second residue in the heme road. The T351 residue in the DsrB subunit is aligned to N393 in the DsrA subunit in the sequence alignment, but structurally they are not in homologous positions (Figure S4). T351 is found at the N-terminal end of a helix in the central interface between heterotetramers in a T351B1-T351B2 interaction, which is the sole interaction at the central interface between heterodimers. The distance between the carbonyl oxygen atoms of the backbone of these two residues is 4.1 Å. This is the third residue of the heme road between the two functional siroheme moieties (Figure 5). Residues N180B1/2 and N393A2/1 are separated by only 10.1 Å, with two intervening aromatic residues, while a single aromatic residue separates N393A2/1 from T351B1/2 (Figure 5D). Thus while the heme road residues are not in direct contact, they are coupled by intervening aromatic side chains.

In support of evolutionary covariance of heme road residues, EVCoupling analysis of the MV2-Eury A subunit also indicated coupling between N369A and L204A, with equally probable residues involved in coupling being R337 and S167. These residues are the structural and sequence homologues of the four residues detected as monomeric FPECs by EVCoupling analysis of the *A. fulgidus* A subunit, three of which constitute the heme road. The third detected FPEC using the *A. fulgidus* B subunit sequence as bait, I350B-R197B (Table 1), implicates K240A and N392A as equally likely to be involved in a residue pair coupling. Residues N392A and I350B are adjacent to two residues in the heme road, N393A and T351B, providing support for their evolutionary coupling. Thus, T351B1-N393A (italicized and underlined in Table 1) are likely to correspond to actual co-varying residues. Given the relatively close proximity of N393B1 and N180B1, 10.1 Å, we hypothesize that these two residues may also be evolutionarily coupled. The aromatic residues linking the heme road residues, Y348B1, F394A2, in *A. fulgidus* are replaced in MV2-Eury by hydrophobic residues L334B1 and I360A2, and F317A1 is replaced by a proline, P166B1, from the B1 rather than A1 subunit.

In the *D. vulgaris* structure, the residues making contact between heterodimers at the central interface are two proline residues (P368B1/B2) that stack on each other (Table 1, Figure 6A), whereas the interacting residues in the MV2-Eury homology model based on the *D. vulgaris* structure are two arginine residues (R337B1/B2) detected in the EVCoupling analysis as possible monomeric FPECs (Table 1, Figure 6B). This central interfacial residue is found to be an asparagine in 33.5% of the sequences and a serine in 7% of the sequences.

**Figure 6.**
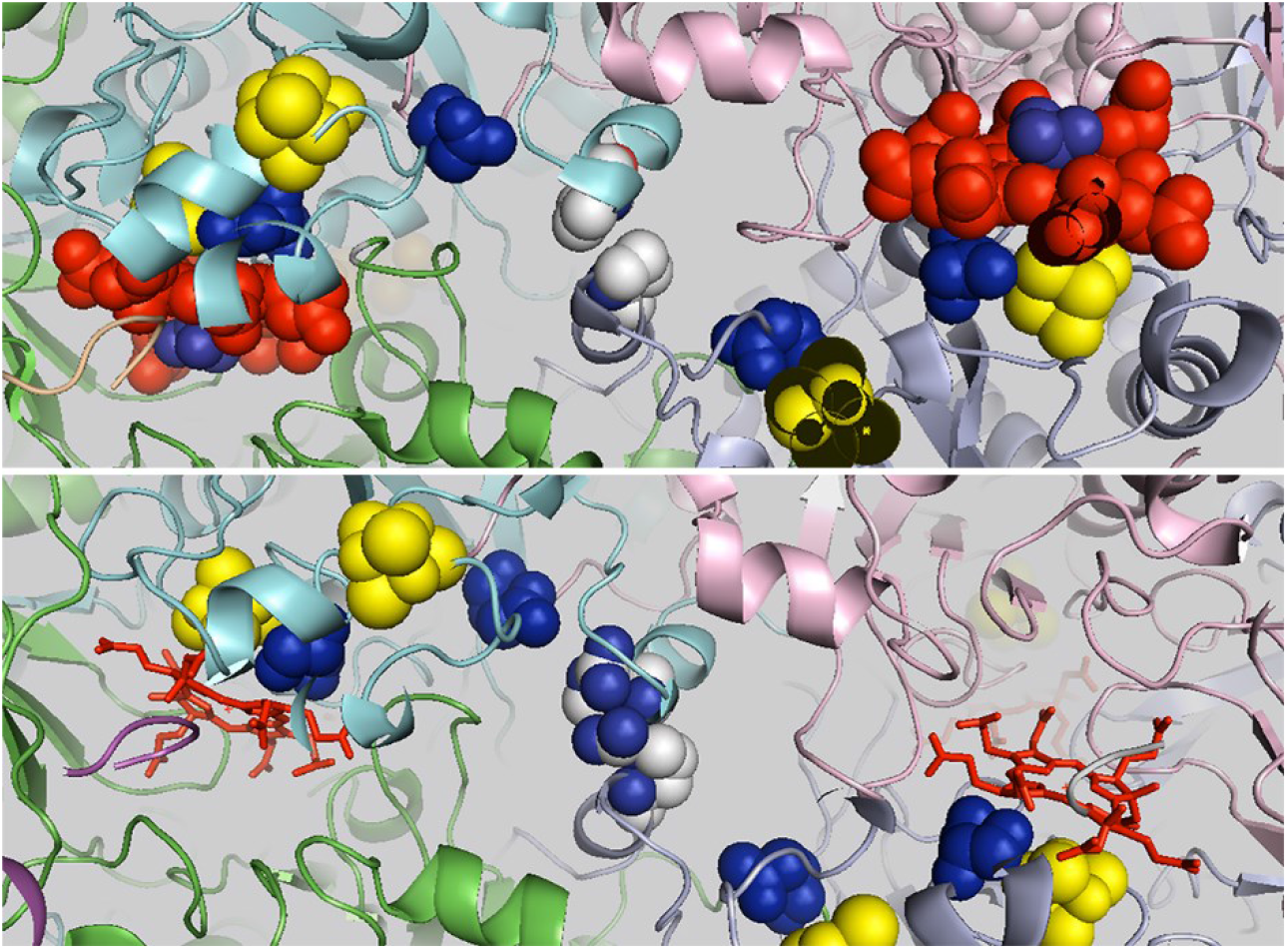
Comparison of the heme road FPEC residues in DsrAB. A). *D. vulgaris* B), the homology model of MV2-Eury. Slab of a zoom from the top views. Chain colors are: A1 (green), B1 (cyan), A2 (light pink), B2 (light blue). Sirohemes (red sticks) and Fe-S centers (yellow) are also shown. FPEC residues (blue spheres) tracing a connection between functional hemes in the *D. vulgaris* structure (A) are from left to right N191B1 (blue spheres), N410A2 (blue spheres), P368B1 (CPK spheres), P368B2 (CPK spheres), N410A1 (blue spheres) and N191B2 (blue spheres) make up the heme road. The corresponding residues in the MV2.Eury model (B) are from left to right S167B1 (blue spheres), N369A2 (blue spheres), R337B1 (CPK spheres), R337B2 (CPK spheres), N369A1 (blue spheres) and S167B2 (blue spheres). Note that the sulfite ion is shown in darer blue bound to the heme in the *D. vulgaris* structure (A).

In 42% of the sequences in the Dsr multiple sequence alignment (MSA) used by the EV Couplings server, the C-terminus is truncated prior to this residue at the central interface, which may correspond to sequencing errors or inclusion of partial sequences in this database. Note that, in the *D. vulgaris* structure, the B1 and B2 helices at the central interface swap positions with respect to the *A. fulgidus* structure (Figure 1). The aromatic residues, F365B1 and Y412A2, in the *D. vulgaris* heme road are switched with respect to Y348B1 and F394A2 in *A. fulgidus*. Like the MV2-Eury heme road, the third aromatic residue in *A. fulgidus*, F317A1 is replaced by a residue from the B1 subunit, I190B1.

Examination of the heme road residues in the structural alignment of the MV2-Eury homology and AF2 models reveals that they are slightly shifted between the two models, with S167 of the B1/2 subunits in the AF2 model exhibiting a steric clash between models with two of the Fe-S centers in the homology model. The two residues constituting the central interface between heterodimers in MV2-Eury B1/2 subunits, R337B1-R337B2, are superimposed at the level of the backbone between the homology and AF2 models, but the side chains point away from each other in the AF2 model, while they interact closely with each other in the homology model (Figure S3). The positioning of the A-subunit C-terminal heme road residue, N369A1/2, in MV2-Eury is quite similar between the homology and AF2 models, despite the very different positioning of the C-terminal helix (Figure 4). The three residues in each heterodimer comprising the heme road on either side of the central interface are not in direct contact, supporting the notion that the co-variance is not uniquely structural in nature. Results of the evolutionary coupling using different subunits from different homologues as bait supports the notion that at least one pair of these residues, T351B1-N393A2 and vice versa in *A. fulgidus*, is truly co-varying. The positioning of N180B1/2 in interaction with the siroheme and relatively close to N393A2/1 is suggestive that this constitutes a second co-varying pair. Based on these observations, we hypothesize that the heme road represents a pathway of cooperative communication between active sites within the DsrAB heterotetramer. The aromatic residues packed between them (Figure 5D) could help modulate information transfer in this putative allosteric route. Alternatively, given the relatively short distances between the heme road residues and the intervening aromatic residues, we cannot rule out that this represents a pathway for electron transfer between heterodimer active sites.

To model the possible dynamic fluctuations that might be associated with this putative allosteric pathway, we carried out Anisotropic Network Modeling (ANM) (40, 41). ANM is based on a Gaussian network that considers protein structures as elastic networks in which the nodes correspond to the Cα atoms, connected by identical spring constants and in which a Kirchhoff matrix is used to represent the topology of internal contacts. ANM has been used to successfully model the known dynamical modes of the allosteric transition in hemoglobin (46). The three most prominent normal modes obtained from the ANM calculation on *A. fulgidus* DsrAB reveal significant motion of one heterodimer with respect to the other, in addition to internal modes within monomers and heterodimers (Figure 7, Movies S1, S2). The red squares along the diagonal in the correlation matrix (Figure S5) indicate motions within each domain of each subunit. Within subunits, individual domains exhibit anti-correlated motions (blue). Off diagonal correlations (red) are observed between subunits within a heterodimer and between heterodimers. For example, motions of domain 1 of the DsrA1 subunit are correlated with the ferredoxin domain of the DsrB1 subunit (Figure S5, yellow circles) and with the domain 1 of the DsrB2 subunit (Figure S5, green circle). The largest motions as evaluated by the calculated B-factors (Figure S6), were observed in the slowest mode for the ferredoxin domains of the DsrA1/2 and DsrB1/2 subunits, and in a region of the DsrB1/2 subunits that is near the structural heme and adjacent to one of the FPEC heme road residues, N180B. The point of contact between the two heterodimers is the FPEC heme road residue, T351. It corresponds to a pivot point (low B-factor) for the most significant modes. Interestingly, the shifts in the backbone observed between the two models of the MV2-Eury sequence, particularly apparent in the ferredoxin domains (Figure 6), mimic the rocking like conformational changes associated with the first two normal modes (Figure 7).

**Figure 7.**
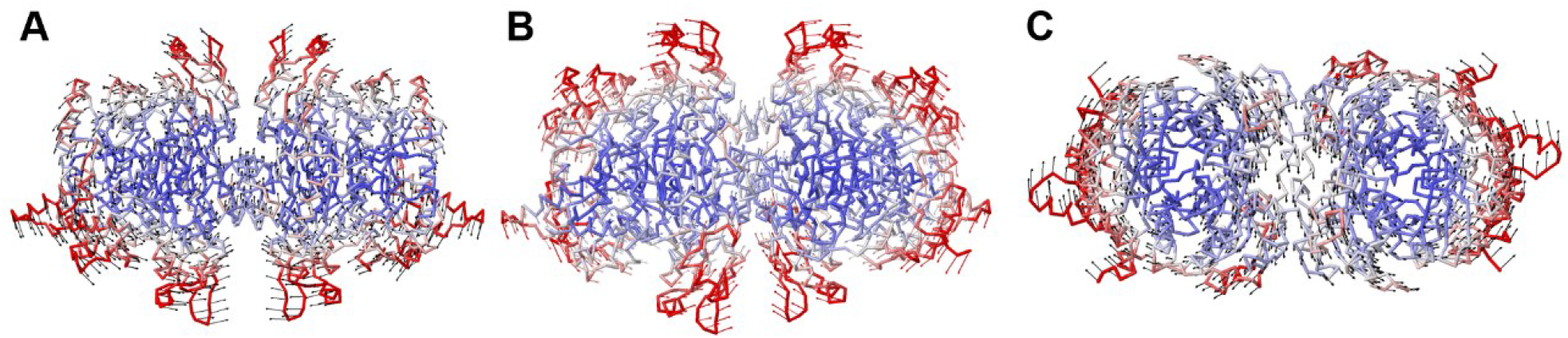
Internal water molecules in the A1B1 heterodimer of the *A. fulgidus* structure of DsrAB. (A) Full heterodimer and (B) Zoom in the siroheme. Chain colors are: A1 (green), B1 (cyan). Sirohemes (red sticks) and Fe-S centers (yellow) are also shown. Internal water molecules are shown as red spheres. The tryptophan residue blocking access to the structural (lower) heme is shown in cpk spheres.

### Inserts and gaps near the structural heme

EVCoupling analysis also revealed another FPEC pair involving residues (D120/K316) near the structural siroheme moiety (in *A. fulgidus*) (bold in Table 2). This pair was identified as an FPEC pair when compared against a contact map of the *D. vulgaris* DsrAB X-ray crystal structure. However, when highlighted on the structure of *A. fulgidus* DsrAB, the residues are in contact. Hence it is a true contact for some members of the protein family, and a false predicted contact for others. Aligning the three structures of DsrAB from *A. fulgidus*, *D. vulgaris*, and the model of MV2-Eury (Figure 8) reveals an antiparallel β-hairpin/loop insert in the *A. fulgidus* DsrB subunit (Figure 8, light pink) which brings aspartate 120B within 7.4 Å of lysine 316B and also makes contact with the end of a helix in the DsrA subunit. This insert includes a tryptophan residue (in CPK color) that blocks access to the structural heme in the *A. fulgidus* structure (see also Figure 3). The MV2-Eury and *D. vulgaris* DsrB subunits (Figure 8, cyan, deep teal) lack this extension. The phenylalanine residue in these sequences, that corresponds to the aspartate in the FPEC pair residue D120B/K316B in *A. fulgidus*, is far (~27 Å) from its co-evolving partner residue (Figure 8, red, raspberry sticks on the right side), thus providing a false positive signal. Interestingly, this insert in the DsrB subunit of *A. fulgidus* (Figure 3, pink) is absent from all other DsrB sequences (Fig. 9), posing the question of why the co-evolutionary coupling probability was high (99%) for this pair.

**Table 2.**
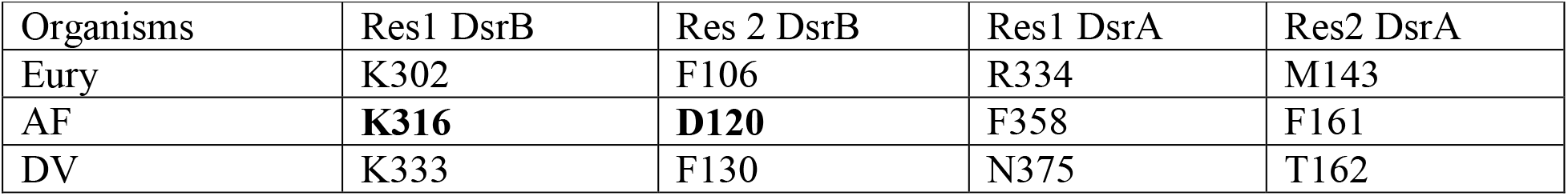
FPEC pairs corresponding to inserts/gaps in the DsrB and DsrA subunits.

**Figure 8.**
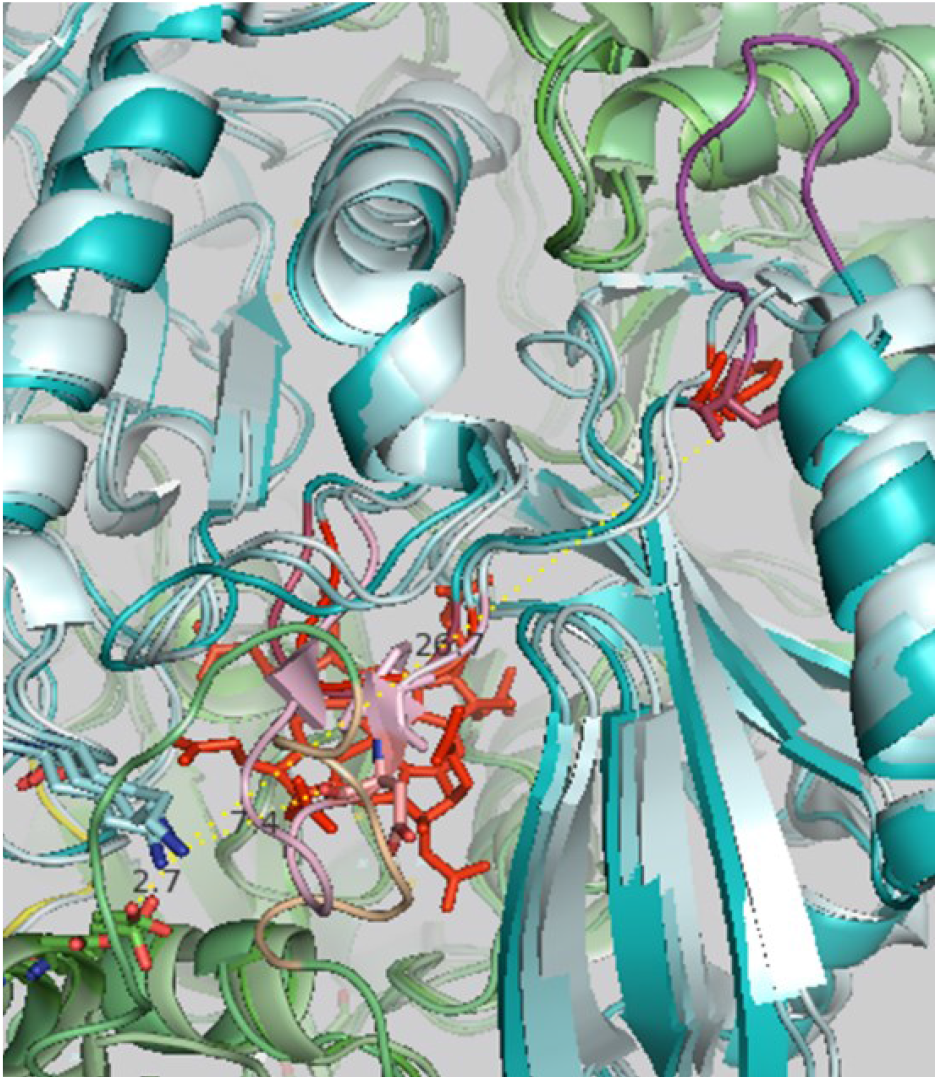
Zoom view of the overlaid structures of DsrAB from *A. fulgidus* (cyan), *D. vulgaris* (dark cyan, and MV2 Eury (pale cyan. The DsrB subunit insert in the *A. fulgidus* structure is in light pink, while the bacterial B loop is magenta. The DsrA subunit insert in the *D. vulgaris* structure is wheat. The rest of the DsrA1 subunits are green. Some of the FPEC residues (K and F/D/F) are indicated in red stick and distances between them (7.4 and 26.7 Å are indicated). The lysine residues of the FPEC form a salt bridge with a glutamate from the DsrA subunit (2.7 Å).

**Figure 9.**
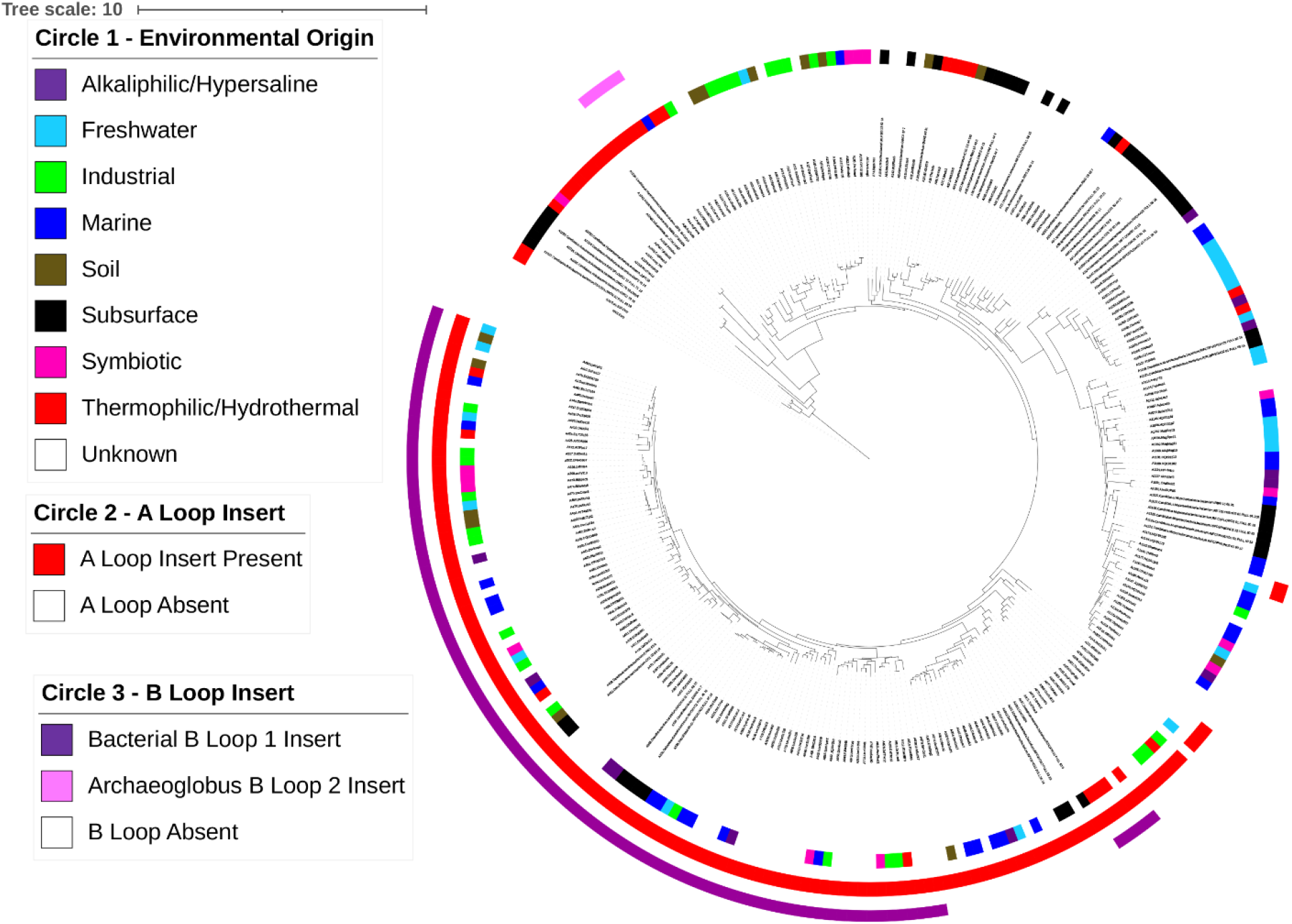
DsrAB phylogenetic reconstruction showing the taxonomic affiliation of Dsr-hosting organisms and the presence of DsrA and DsrB structural inserts. The Maximum Likelihood reconstruction was conducted on a concatenated alignment of DsrA and DsrB subunits from a curated database comprising the previously recognized primary homolog groups. Scale bar shows the expected substitutions per site. Taxonomic classification is given based on information in the database (Muller et al. 2015) or by >80% amino acid homology to Dsr from cultivars/genomes with taxonomic annotation. DsrA or DsrB inserts were identified based on structural characterizations and subsequent identification within DsrA or DsrB alignments.

The *D. vulgaris* sequence exhibits another insert in the DsrB subunit (Figure 8, magenta), just prior to that observed in the *A. fulgidus* sequence that makes contact with a different helix in the DsrA subunit. This insert is missing in all early-branching DsrAB (Figure 9, outer circle, purple) and is present in nearly all Deltaproteobacteria, in addition to a few other lineages with evidence for HGT of DsrAB from Deltaproteobacteria to other lineages (i.e., Thermodesulfobacteria and some Firmicutes (23); Figure 9). Thus, this adaptation appears to have evolved in the Deltaproteobacteria and has been retained throughout this lineage, in addition to maintenance in other lineages following HGT. These results suggest that these adaptations were a consequence of the ecophysiological lifestyles of late-evolving Deltaproteobacteria.

## Discussion

The diversity of SRO and their DsrAB enzymes has been greatly expanded through recent cultivation- and cultivation-independent approaches (11, 23, 24). Together these studies suggest that Dsr likely emerged to catalyze SO_3_^2-^ reduction and then diversified (through recruitment of APS and Sat) to catalyze SO_4_^2-^ reduction and ultimately HS-oxidation (9–12). The phylogenetic studies conducted herein suggest that model bacterial SROs implicated as major players in contemporary biogeochemical S cycling (e.g., Deltaproteobacteria and Firmicutes) evolved comparatively recently whereas early evolving archaeal SROs (and a few taxonomically patchy bacterial genera) tend to be restricted to hydrothermal or more nutrient-limited extreme environments (11) where oxidant limitation is likely pervasive (47). While the ecological drivers of the evolution of SROs (via Dsr phylogeny) is obscured in the present study by limited corresponding metadata (e.g., cardinal growth parameters, geochemistry) associated with these organisms or the environments from where they were recovered, the broad differences in habitats of early evolving and later evolving SROs suggests that the ecology of Dsr-harboring SROs has evolved over time. Yet, it remains unclear if the structure and thus, functional mechanisms of Dsr has also evolved during its evolutionary history.

Despite the collective abovementioned observations indicating SRO (and thus Dsr) diversification across gradients in temperature, pH, salinity, and pressure, amongst other variables, the phylogenetic studies conducted herein and elsewhere document a general pattern of vertical inheritance and a high degree of overall primary sequence conservation across all DsrAB (11, 23, 24). Consistent with these findings, available structures and our structural models of selected Dsr enzymes that represent much of the known sequence diversity of Dsr generated herein reveal a high degree of structural conservation. Dsr forms a heterotetrameric structure comprising two heterodimers of DsrA and DsrB, the latter of which arose from an ancestral gene duplication (22, 48). The high degree of structural conservation, including at the inferred quaternary level, suggests that all extant lineages of SROs settled on this Dsr structural configuration prior to the radiation of SROs.

The evolutionary co-variance in the extensive inferred conservation in the structure of Dsr revealed a group of three residues in each heterodimer that form a pathway between the two active site sirohemes. Based on the homologous monomeric FPECs from both the *A. fulgidus* and MV2-Eury A subunit queries, and a second set of monomeric FPECs from the *A. fulgidus* B subunit query adjacent to two of the three residues, we conclude that the N393A2/1-T351B1/2 (*A. fulgidus*) pair is truly coupled evolutionarily. Given the position of N180B1/2 interacting with the active site heme and the Fe-S center, and its relatively close proximity to the N393A2/1 residue, we hypothesize that N180B1/2-N393A2/1 are also evolutionarily coupled. Based on structural link formed by these residues between the two active sites, we hypothesize that, beyond simply stabilizing contacts between heterodimers, the pathway traced by these residues could correspond to an ancient allosteric pathway (i.e., the “heme road”) that could have provided the advantage of allosteric control when the heterodimer to heterotetramer transition occurred. While these heme road residues are not in direct contact, their interactions are mediated by only one or two aromatic residues. Consistent with the existence of an allosteric pathway are the hinge motions predicted by ANM. Inter-domain motions in multi-chain protein complexes are an increasingly appreciated aspect of their dynamic structures and thermodynamics, with the potential to modulate their functions (49–51). Our Gaussian Network normal mode analysis (40, 41, 46) indicates the possibility of intersubunit “rocking” in the structure of the DsrAB complex, like that observed for other dimeric complexes (49). These motions could in principle be frozen out by crystallization of multiple members of the DsrAB family, as also observed across crystal structures of dimeric influenza NS1 protein domains (49). In further support of such interfacial dynamics in DsrAB complexes, which could also relate to the proposed intersubunit allostery, is the lower quality of the electron density maps noted by the authors of the *A. fulgidus* structure between the A2B2, as opposed to the A1B1, heterodimers (20). This difference in quality could reflect an asymmetry in the structures of the two heterodimers due to allosteric interactions.

The proposed heme road allosteric pathway could serve to allow for communication between the two active sites, one in each heterodimer, during the delivery of 2 e^-^ from DsrC to one of the active sites. Such an interaction could inhibit DsrC binding and electron injection into one heterodimer, while the other is active. This is the case for negative cooperativity in the function of the Mo-Fe nitrogenase heterotetramer (29), which exhibits similar rocking normal modes as DsrAB. Examples of statistically detected evolutionarily conserved pathways of energetic coupling within proteins have been reported for several proteins, including PDZ domains, GPCRs, chymotrypsin, lectin and hemoglobin (27, 52–55). We emphasize the importance of experimental validation of putative allosteric networks revealed by statistical analysis of sequence co-evolution. Such validation can be accomplished by combinations of approaches including H/D exchange mass spectrometry (56) and/or NMR (57), and spectroscopic approaches coupled with mutagenesis (58). We note that given the distances, we cannot rule out that the heme road corresponds to a pathway for electron transfer between active sites, although it is difficult to rationalize its utility.

While this proposed pathway warrants experimental scrutiny, the co-evolution at the positions putatively involved also indicates that this pathway was likely established prior to the radiation of all DsrAB, although the precise residues involved in these interactions vary across Dsr enzyme types. Consequently, the presence of a slightly modified allosteric pathway may have allowed fine-tuning of Dsr activity in the context of different physiological backgrounds, including those that operate under chronic energy (dissimilatory e^-^ shuttling) stress imposed by extreme conditions (e.g., temperature, pH and pressure) (47) where the kinetics of HSO_3_^-^ reduction and thus growth of SRO are likely to be much slower. The combination of the evolutionary coupling analysis and ANM dynamic “normal mode” calculations provide an intriguing set of residues to probe for their involvement in structure-function relationships in future experiments.

In addition to the aforementioned putative allosteric pathway, it is possible that the presence of the W119 residue in *A. fulgidus* Dsr and the loss of the iron in the sirohydrochlorin moieties in *D. vulgaris* Dsr represent two separate mechanisms to limit function to the “top” siroheme in each heterodimer, adjacent to the binding site of the DsrC putative electron carrying subunit. Evolution of the gaps and inserts in the region of the structural hemes may have contributed to their loss of function. Clearly, detailed comparative enzymatic assays will be required to demonstrate the structural, as opposed to functional, role of the “bottom” siroheme”.

Collectively, the combination of phylogenetic and structural bioinformatics studies of Dsr conducted herein point to the need for comparative experimental studies to test the present hypotheses in order to fully understand the functional differences encompassed within the diversity of these enzymes. This is particularly true for the two most-basal branching lineages of Dsr from SRO that are most reminiscent of the ancestral Dsr enzymes that likely shaped sulfur and carbon biogeochemical cycles on early Earth and that continue to shape these cycles in thermal environments. The vast majority of these taxa (inclusive of MV2-Eury) are either known only from cultivation-independent environmental genomics studies or have very-limited cultivation information. Thus, future efforts should first be made to domesticate these SROs and optimize cultivation conditions to enable more thorough investigations of their physiology, ecology, and enzymology. In particular, these efforts should focus on SROs that are physiologically unlike canonical SROs that conduct SO_4_^2-^ reduction but that rather are limited to SO_3_^2-^ reduction. Such investigations would enable a better understanding of the evolution of Dsr as it transitioned from early SO_3_^2-^ respiring organisms to the SO_4_^2-^ reducers that are widespread in anoxic environments on Earth today.

## Acknowledgments

We thank Drs. Kelly Brock and Chris Sander for assistance with sequence co-variance analysis using EvFold. DRC and ESB acknowledge support from the National Science Foundation (EAR-1820658) and the NASA EPSCoR program (MT-19-EPSCoR-0020). GTM acknowledges support by NIH Grant R35-GM141818.

