## Supplemental Figures for "Structural Evolution of the Ancient Enzyme, Dissimilatory Sulfite Reductase"

and Catherine A. Royer*

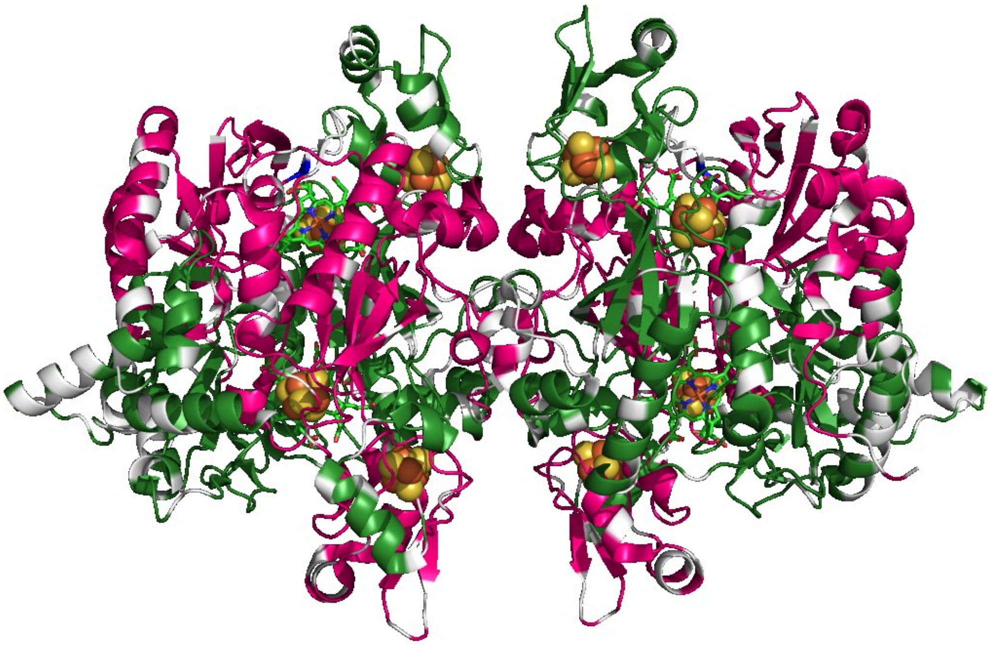

**Figure S1**. Amino acid residue positions for substitutions between DsrAB from *A. fulgidus* and *D. vulgaris*. Substituted residues are shown in grey.

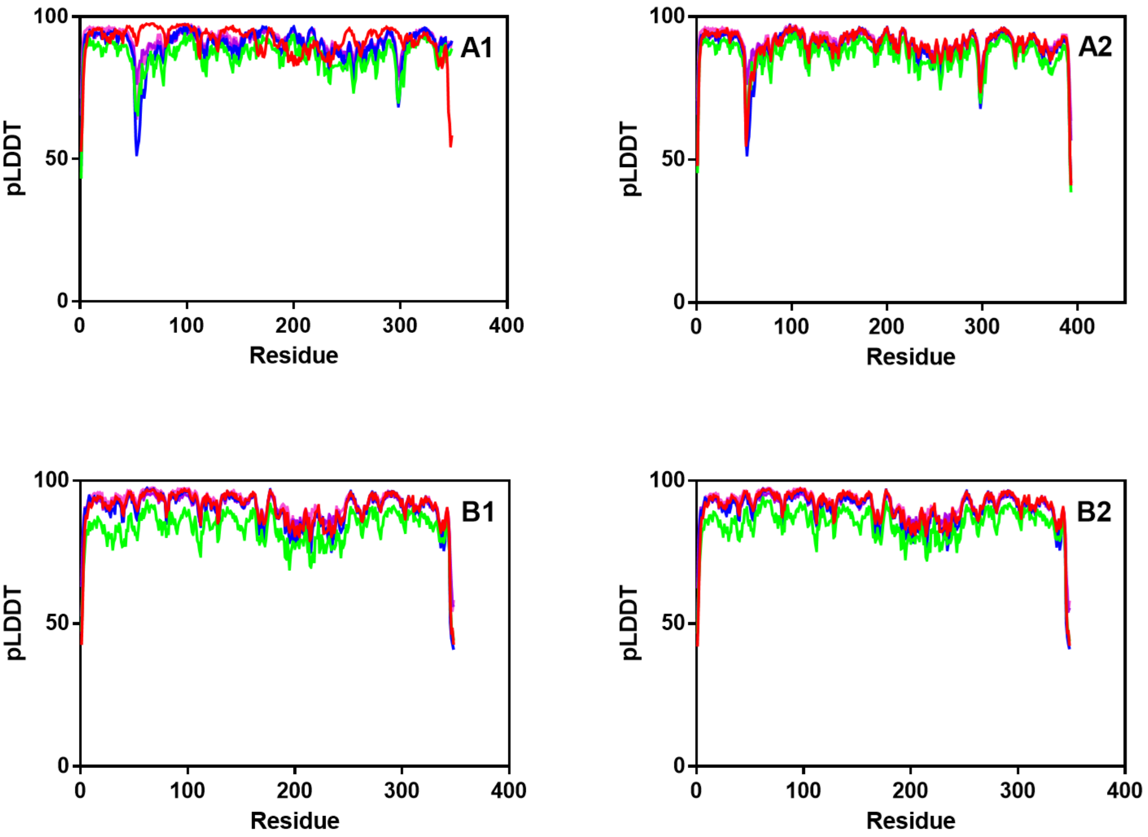


**Figure S2**. Results of structural modeling the heterotetrameric complex of the MV2-Eury sequence using AlphaFold2 (AF2) Advanced Multimer. pLDDT scores represent the quality of the model, with scores near 100 corresponding to very high confidence. pLDDT scores for each of the four subunits of the heterotetramer are shown for the three models obtained from AF2, as indicated. Model 1-red, Model 2-green, Model 3-blue, Model 4-violet, Model 5-pink. Model 3 gave the highest pLDDT score (0.8340) across all subunits, followed by Model 1 (0.8302), Model 4 (0.8277), Model 5 (0.8271) and Model 2 (0.7656).

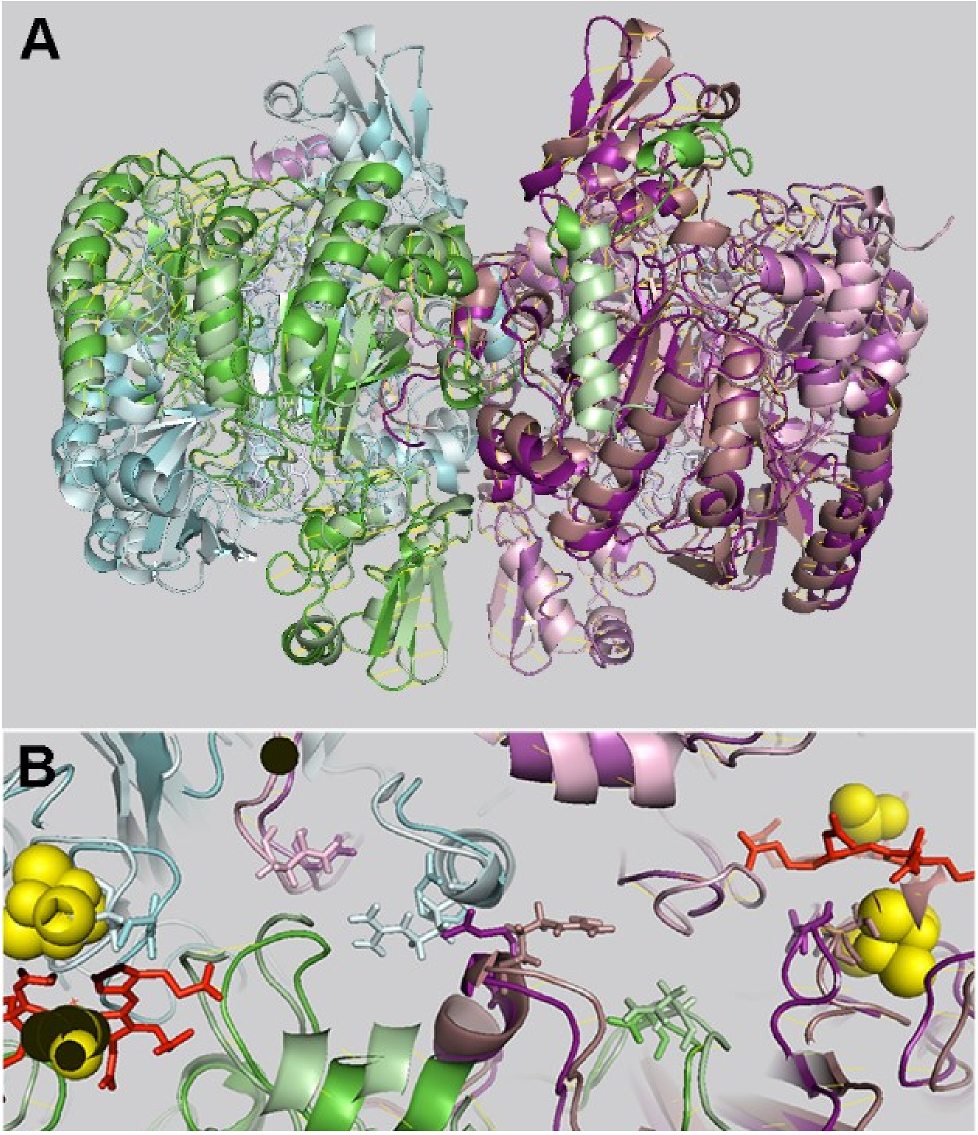


**Figure S3. Alignment of the AlphaFold2 MV2-Eury with the homology model.** A) The homology model subunits are colored A1-green, B1-cyan, A2-magenta, B2-deep purple. Those for the AF2 model 1 are colored as follows: A1-light green, B1-light cyan, A2-light pink, B2-dirty violet. Global structural overlay of the two models, homology and AF2. The two heterodimers are more rotated with respect to each other around the central access at the heterodimer interface. B) Overlay of the heme road residues from the homology and AF2 models of MV2-Eury The B1/2 interfacial residue, R337, from the MV2-Eury homology and AF2 models (B1-cyan, light cyan, B2-dark purple, dirty violet, respectively) shows the two side chains from the B1 and B2 subunits facing in opposite directions in the AF2 model, whereas in the homology model, they interact (also see Figure 5). The rest of the heme road residues, B1/2: S167 near the Fe-S center and A1/2:N369 situated in between the other two, are shifted in the two models with respect to each other due to the rotation of the heterodimers around the central axis.

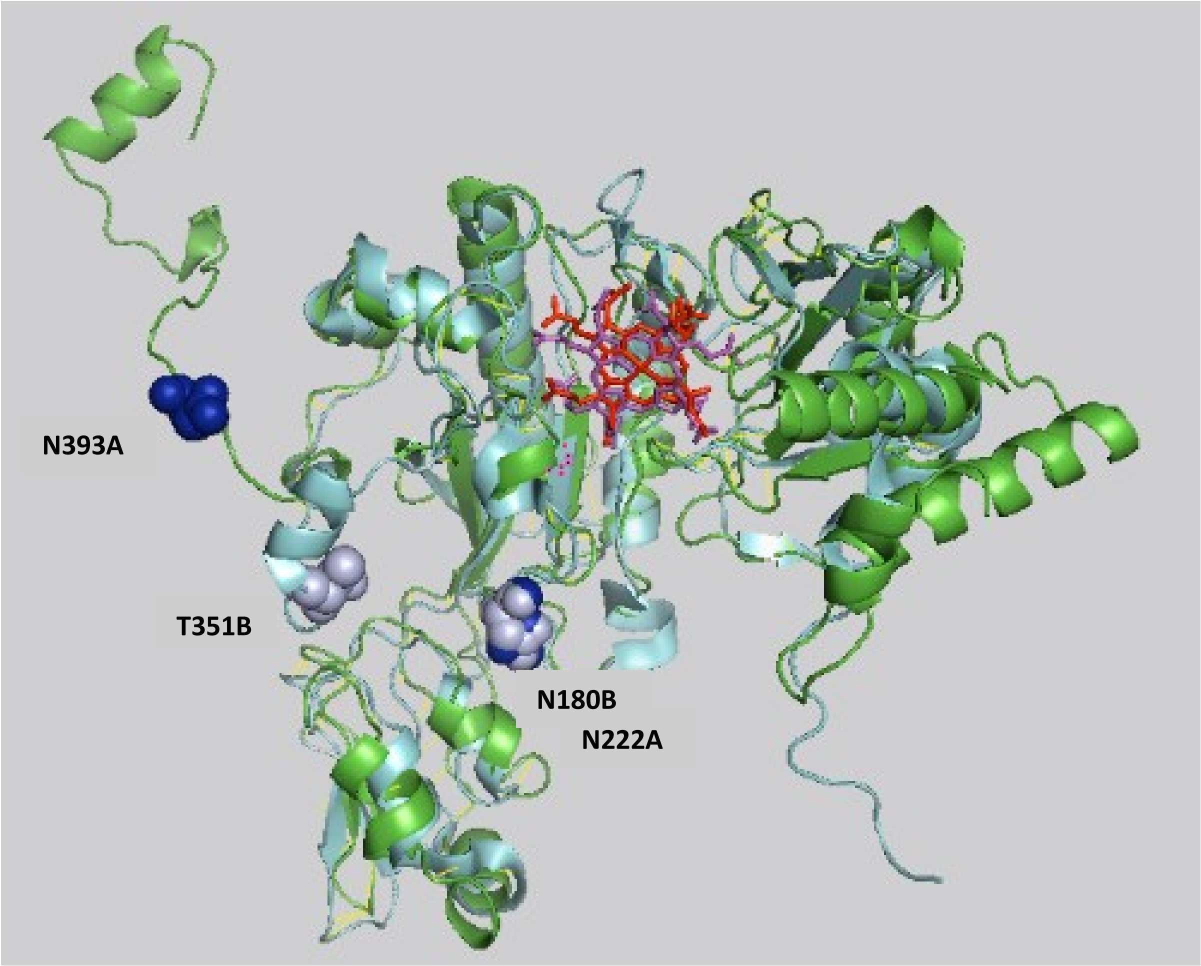


**Figure S4. Alignment of the structures of the DsrA and DsrB subunits from *A. fulgidus*.** The DsrA subunit is in green and the DsrB subunit in cyan. The DsrA subunit heme is in red and the DsrB subunit heme is in magenta. The DsrA subunit heme road residues (N222 and N393) are in dark blue and the DsrB subunit heme road residues (N180 and T351) are in light blue. It can be seen that while N180B and N222A occupy nearly identical positions in the aligned subunit structure above, in the heterotetramer, the N180B residue is actually near the magenta (active) siroheme of the DsrA subunit, while the N222A residue is proximal to the inactive, structural heme of the DsrB subunit

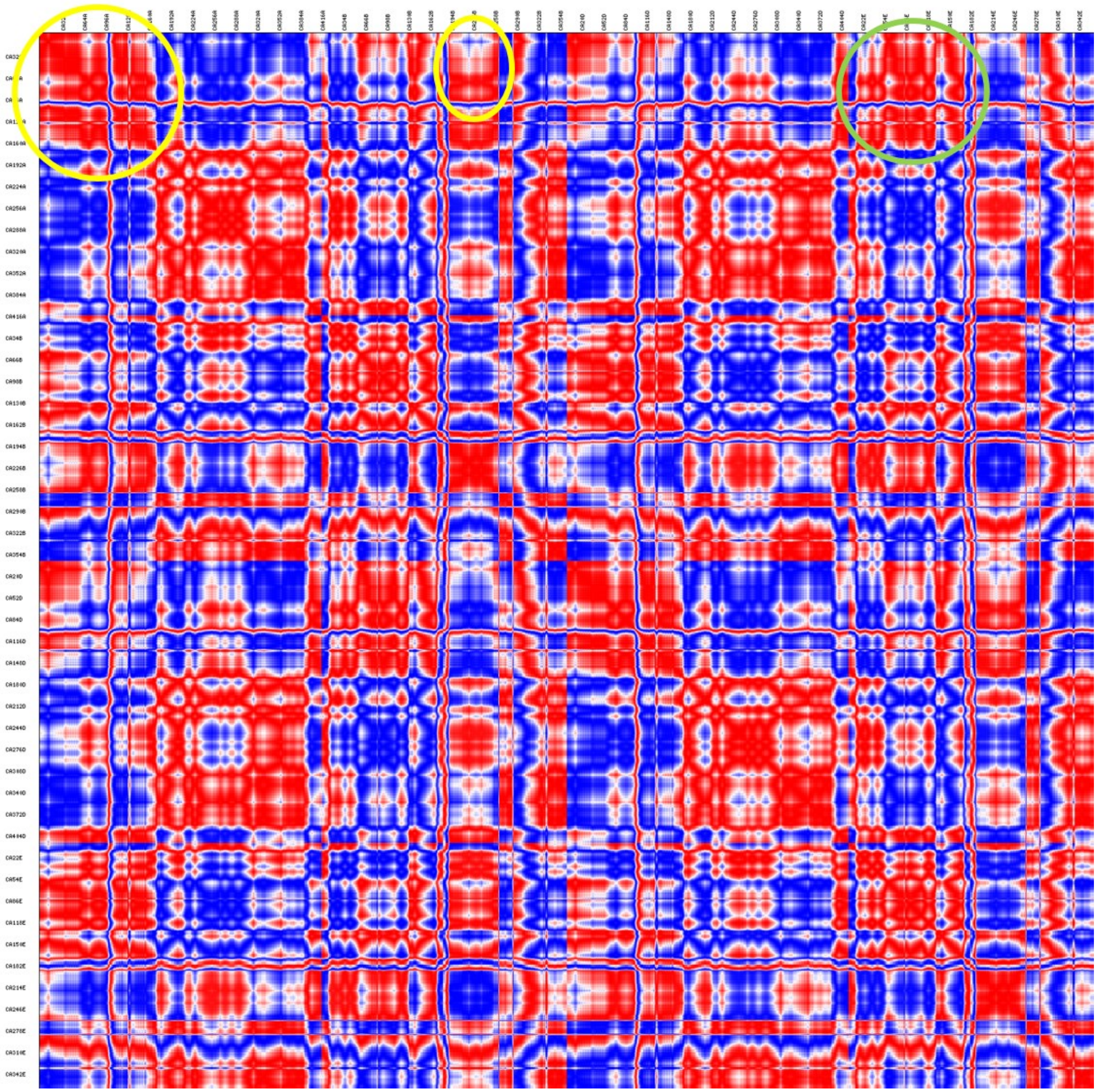

**Figure S5. Correlation matrix for motions in the DsrAB heterotetramer.** Motions were calculated using Anisotropic Network Modeling as described in the text. Red values are positively correlated motions, while blue corresponds to negatively correlated motions.

**Chain A**

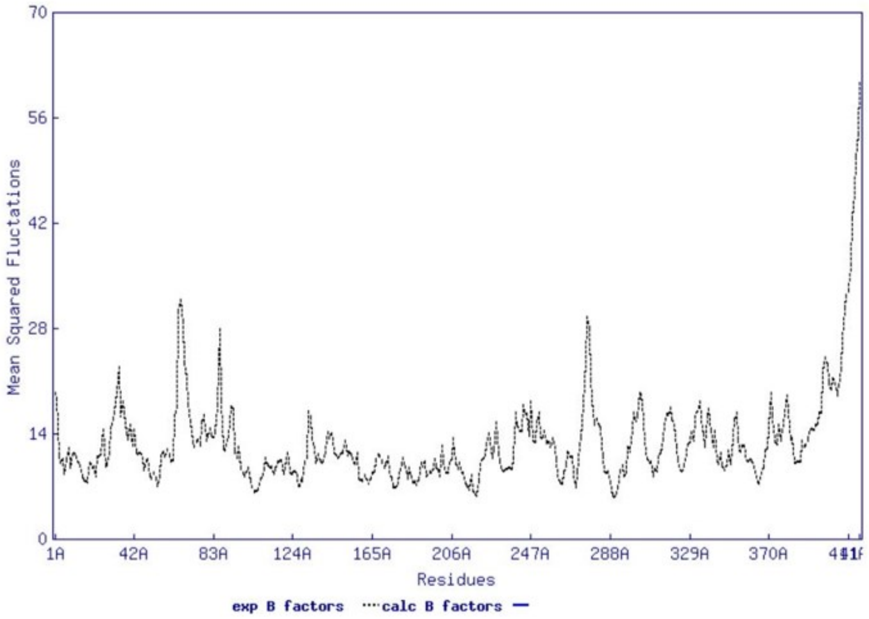


**Chain B**

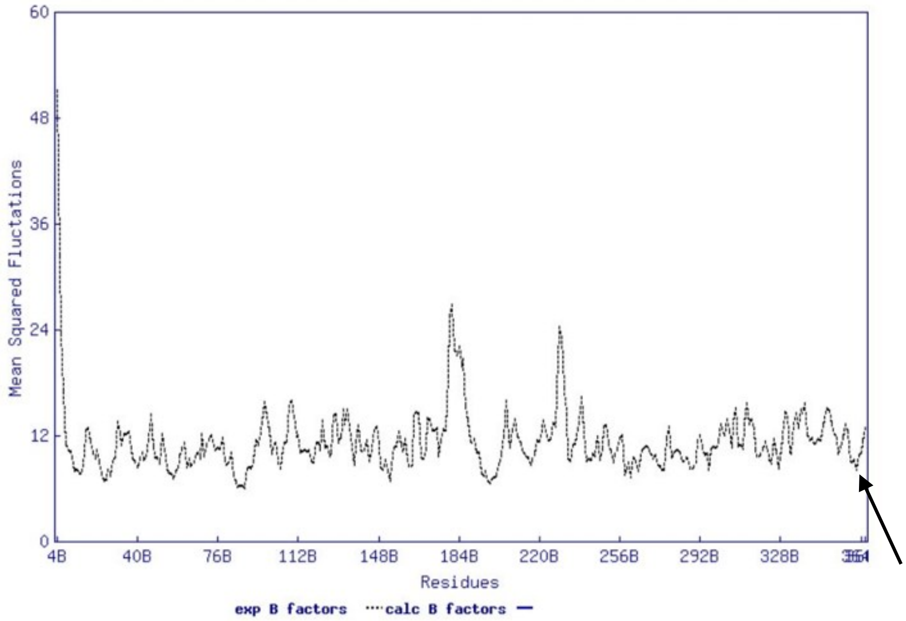


**Figure S6. "B-factors" calculated for the residues of the DsrA and DsrB chains of the heterotetramer.** Values were obtained from the Anisotropic Network Modeling procedure as described in the text. The first minimum in chain B beginning at the C-terminus of the chain (arrow) corresponds to residue BT351 in the *A. fulgidus* sequence. Low B-values are consistent with its position as a central pivot point in the dynamics motions of one heterodimer with respect to the other.
